# Metabolic collapse as a mechanism of developmental regression: convergent evidence from Kleefstra syndrome [^18^F]FDG-PET/CT imaging and *Drosophila* modelling

**DOI:** 10.64898/2026.08.28.747020

**Authors:** Spencer G. Jones, Arianne Bouman, Nicholas Raun, Evelien A.J. van Genugten, Isabel Martínez-Blázquez, Franziska Kampshoff, Janine Doorduin, Joyce Geelen, Hilgo Bruining, Karlijn Vermeulen-Kalk, Stéphanie Miot, David Geneviève, Erik H.J.G. Aarntzen, Mireia Coll-Tané, Tjitske Kleefstra, Annette Schenck

**Author notes:** These authors contributed equally (co-first authors) (Spencer G. Jones, Arianne Bouman). These authors contributed equally (co-last authors) (Tjitske Kleefstra, Annette Schenck).

## Abstract

Developmental regression is a severe but poorly understood complication of several neurodevelopmental disorders. In Kleefstra syndrome (KLEFS1), caused by *EHMT1* haploinsufficiency, regression often emerges during adolescence or early adulthood and is frequently preceded by marked sleep disturbance. Experimental work implicating *EHMT1/G9a* in metabolic regulation and stress responses raises the possibility that impaired metabolic resilience contributes to this vulnerability. Here, we aimed to investigate whether altered glucose metabolism is a feature of KLEFS1 and whether it relates to clinical variability, including regression. Through [^18^F]FDG-PET/CT, individuals with KLEFS1 who had experienced regression (n=4) exhibited a hypometabolic brain profile, whereas one individual who had not experienced regression showed globally elevated metabolic activity. In parallel, *G9a* mutant flies exhibited increased baseline metabolic rate and neuronal ATP levels together with sleep fragmentation resembling the clinical phenotype. Providing flies with oxidative stress to model KLEFS1 regression further exacerbated sleep disruption and was associated with a reduction in metabolic output. Importantly, adult high sugar feeding in flies prevented oxidative stress-induced worsening of sleep and maintained metabolic stability under challenge. Together, these findings suggest that regression in KLEFS1 and associated sleep disturbances are linked to underlying metabolic vulnerability and impaired maintenance of energy homeostasis under stress.

## Introduction

Developmental regression is a severe and poorly understood complication associated with several neurodevelopmental disorders (NDDs), including autism spectrum disorder, Phelan-McDermid syndrome, Rett syndrome, Down syndrome, Fragile X syndrome, 22q11.2 deletion syndrome, and Kleefstra syndrome^1^. Characterized by the loss of previously acquired developmental or adaptive skills, regression can profoundly affect cognition, communication, motor function, and daily functioning, however, the biological mechanisms underlying susceptibility remain largely unknown.

Kleefstra syndrome (KLEFS1; OMIM #610253) is a rare neurodevelopmental disorder caused by haploinsufficiency of *EHMT1*, which encodes euchromatic histone methyltransferase 1, a key regulator of chromatin structure and gene expression^2–5^. Individuals with KLEFS1 present with intellectual disability, developmental delay, childhood hypotonia and distinctive craniofacial morphology^5–7^. They also exhibit a high prevalence of comorbidities, including autism spectrum disorder (ASD), childhood-onset obesity (∼60%), and sleep disturbances^6,8–11^. Developmental regression is a clinically significant complication of KLEFS1, affecting 11–50% of individuals across cohorts^6,10,12,13^, and typically presents as catastrophic regressive episodes during adolescence or early adulthood^14–17^.

Regressive episodes in KLEFS1 have been associated with physiological or psychological stressors, including infection, puberty-associated hormonal changes, and major life events, suggesting that diverse environmental challenges may precipitate regression in individuals with underlying vulnerability^11,12,18^. Severe sleep disturbances frequently precede or accompany regressive episodes and are recognized clinically as an early indicator of impending regression^14,19,20^. Given the essential role of sleep in cognitive function^21^, sleep disruption may further contribute to adaptive skill loss. Although sleep interventions can improve behavioural outcomes^22^, the mechanisms linking sleep disruption to regression and other manifestations of KLEFS1 remain poorly understood.

Metabolic control has emerged as a critical function of *EHMT1*, and preclinical studies in mice and flies show that loss of *EHMT1* or its orthologs (*GLP*/*G9a*) can impair both the development of energy-regulating tissues and stress-responsive energy homeostasis^23–27^. In mice, knockout of *GLP* in differentiating brown adipocytes impairs brown fat development and promotes obesity and insulin resistance^26^. In *Drosophila*, *G9a* mutants present dramatically elevated energy stores, but when exposed to stressors such as oxidative stress (OS) or infection, glycogen stores are rapidly depleted while lipid stores inefficiently mobilize, resulting in premature death^23,24,28^. These findings support a critical role for *EHMT1* in buffering metabolic responses to stress, maintaining energy homeostasis, and protecting neuronal and systemic function. Building on this, it has been hypothesized that individuals with KLEFS1 may also experience reduced accessibility of fat stores as an energy source, contributing to observed obesity^8^, and that stress induced metabolic catastrophe may lead to systemic glucose depletion that lowers brain glucose availability, potentially triggering developmental regression, sleep disturbances, and cognitive decline^24,29^. In line with this model, at least one KLEFS1 patient experienced a pneumonia infection preceding the onset of severe sleep disturbances and regression^14^, and similar stressors, including infection and injuries, have been reported to trigger delirium followed by impaired glucose utilization in neurodegenerative disorders^30^.

To date, no clinical studies have investigated the pathophysiological mechanisms underlying metabolic dysfunction potentially contributing to regression in KLEFS1, yet insight into these pathways is critical for understanding neurodevelopmental vulnerability. In the current study, we set out to assess glucose metabolism in individuals with KLEFS1 using [^18^F]fluorodeoxyglucose positron emission tomography/computed tomography ([^18^F]FDG-PET/CT), focusing on the brain and peripheral tissues, including the liver, spleen, blood pool, bone marrow, muscle, and adipose tissue. To complement these clinical findings and generate hypotheses about potential mechanisms, we used an established *Drosophila* G9a loss-of-function model which exhibits baseline sleep disturbances resembling the clinical phenotype^29^. Building on prior observations that energy availability is critical for coping with environmental stress^23^, we used OS-induced exacerbated sleep disruption as a readout relevant to human regression, enabling us to explore how metabolic fragility might contribute to regressive phenotypes. By combining exploratory [^18^F]FDG-PET/CT imaging with complementary studies in a *Drosophila* G9a model, we investigated whether convergent evidence supports altered metabolic regulation as a potential contributor to regression vulnerability in KLEFS1.

## Results

### Clinical and demographic characteristics of the KLEFS1 cohort

To better define the clinical and metabolic profiles associated with KLEFS1, we examined a cohort of five adult individuals with molecularly confirmed diagnoses using [^18^F]FDG-PET/CT. Participants 1-3 received whole body [^18^F]FDG-PET/CT scans (toes to skull) under general anesthesia at Radboudumc, while participants 4-5 received brain [^18^F]FDG-PET/CT scans in France, with Participant 4 having undergone two scans approximately one year apart. Demographic data of participants is shown in **Table 1**. All individuals had intellectual disability, and three (Participants 1, 4 and 5) had a clinical diagnosis of autism spectrum disorder. A regressive episode occurred in four participants (1, 3, 4, and 5), while Participant 2 had no history of regression; regressive episodes, which exceeds the traditional DSM classification, were defined as an absolute decline in functioning in at least one of the adaptive behavior domains of practical, conceptual, or social-emotional skills in the individual, which, when left untreated, would last at least several months^1,11,17^. Psychiatric symptoms may accompany regressive episodes but are not required for their definition, these are variable between individuals. Psychosis accompanied the regression in two cases (Participants 1 and 3) and was treated with olanzapine, which may influence brain glucose metabolism^31^. Participant 5 had active *grand mal* epilepsy experiencing approximately one generalized tonic-clonic seizure per month, while Participant 4 had a past history of seizures no longer requiring antiepileptic treatment. Sleep disturbances were noted in four individuals (Participants 1, 3, 4 and 5), and recurrent infections were reported in two (1 and 2). Metabolic laboratory evaluation at the time of scanning was available for Participants 1-5, all of whom showed glucose values within the range of normal glucose. Participant 2 exhibited elevated ketone bodies and insulin with normal glucose (HOMA-IR 5.0), indicative of insulin resistance; Participant 1 showed folic acid deficiency, while no additional metabolic abnormalities were reported for Participants 3-4. Overweight or obesity was present in three participants (1-3), with BMI values ranging from 25.6 to 31.9.

**Table 1.** Demographic data of participants.

| Table 1. Demographic data of participants |  |  |  |  |  |  |  |  |  |  |  |  |  |  |  |  |
| --- | --- | --- | --- | --- | --- | --- | --- | --- | --- | --- | --- | --- | --- | --- | --- | --- |
| # | Age (yr.m) | Sex | MBq | Genetic variant (GRCh38) | ID | ASD | Regression | Psychosis | Depression | Anxiety | Epilepsy | Sleep disturbance | Recurrent infections | Metabolic features | Body mass Index (BMI) | Medication |
| 1 | 26.6 | F | 228.5 | c.1968dup; p.Gln657AlafsTer41 | M | Y | Y | Y | N | Y | N | Y | Y | Folic acid deficiency | 26.4 | Propofol 60mg<br>Olanzapine 12.5mg |
| 2 | 21.8 | F | 331 | c.2587C>T; p.Gln863Ter | M | N | N | N | N | N | N | N | Y | Insulin resistance, Ketosis at PET/CT scan | 31.9 | Propofol 80mg, Alfentanil 0.5mg |
| 3 | 29.5 | F | 233 | chr9:g.(137501727_137505613)_qterdel | S | N | Y | Y | Y | Y | N | Y | N | N | 25.6 | Remimazolam 7mg<br>Citalopram 10mg<br>Olanzapine 5mg |
| 4 | 38 | F | 125 (Scan #1), 115 (Scan #2) | c.2712+1G>A | M/ S | Y | Y | N | N | Y | N (past history) | Y | N | N | 21.7 | Alprazolam 0.75mg |
| 5 | 21.4 | M | 118 | c.2566C>T, p.(Gln856Ter) | S | Y | Y | N | N | Y | Y | Y | N | N | 15.9 | Lamotrigine 200mg<br>Clobazam 30mg<br>Lacosamide 200mg<br>Perampanel 2mg |

MBq: MegaBecquerel [<sup>18</sup>F]FDG injected before scan. Y: Yes, N: No. ID: intellectual disability; M: Moderate, S: Severe. ASD: Autism
Spectrum Disorder

### Brain [^18^F]FDG metabolic profiles in KLEFS1

For our KLEFS1 cohort and healthy controls (n=5), four different brain regions were evaluated for [^18^F]FDG uptake: the frontal lobe, temporal lobe, basal ganglia, and cerebellum, with mean standardized uptake values (SUVs) summarized in **Table S1**. Two distinct metabolic patterns emerged. Participant 2 demonstrated global brain hypermetabolism, with elevated SUVs across all brain regions relative to controls **(Fig. 1A,B)**. In contrast, Participants 1 and 4 showed widespread hypometabolism affecting all examined regions. For Participant 4, this pattern was consistent across two brain [^18^F]FDG-PET/CT scans acquired approximately one year apart, with hypometabolism observed in all four regions in both scans. Participant 3 displayed selective frontal hypometabolism, while temporal, basal ganglia and cerebellar uptake remained generally within the normal range. Participant 5 displayed frontal hypometabolism and mildly elevated basal ganglia metabolism, with temporal and cerebellar uptake within the normal range. Overall, these findings indicate that Participant 2, who did not experience clinical regression, exhibited a globally elevated brain metabolic profile, whereas those who experienced regression showed reduced or region-specific decreases in brain metabolism.

**Figure 1.**
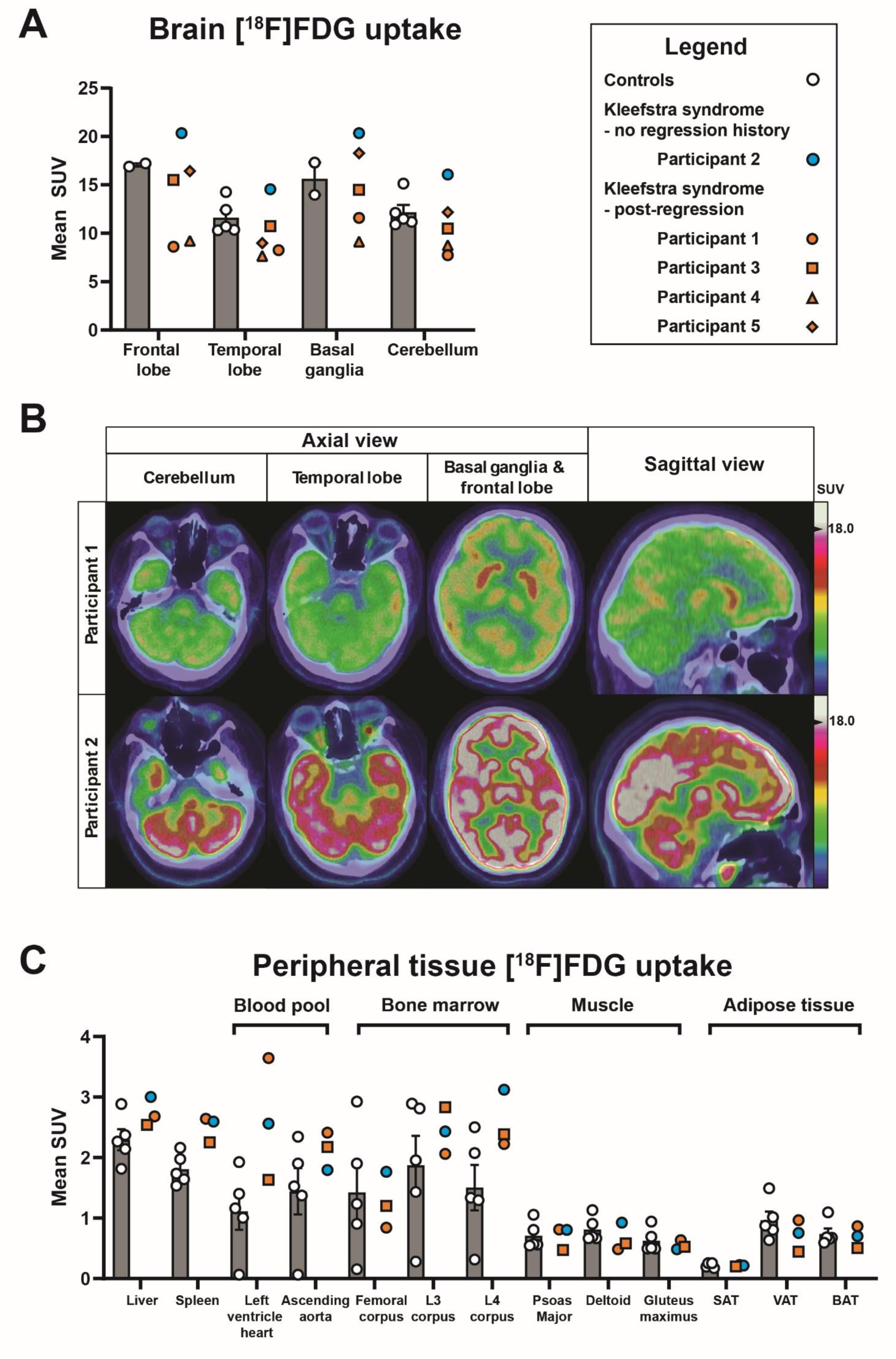
Brain and peripheral tissue [^18^F]FDG uptake in individuals with KLEFS1 and controls. (A) Mean standardized uptake values (SUVs) for cerebral [^18^F]FDG uptake in the frontal lobe, temporal lobe, basal ganglia, and cerebellum. (B) Representative axial and sagittal [^18^F]FDG-PET/CT images of Participants 1 and 2, showing [^18^F]FDG uptake in the cerebellum, temporal lobe, basal ganglia, and frontal lobe. (C) Mean SUVs for peripheral tissue [^18^F]FDG uptake in the liver, spleen, blood pool (left ventricle and ascending aorta), bone marrow (femoral corpus, L3 and L4 vertebral bodies), skeletal muscle (psoas major, deltoid, and gluteus maximus), and adipose tissue, including subcutaneous adipose tissue (SAT), visceral adipose tissue (VAT), and brown adipose tissue (BAT). The y-axis shows the mean SUV, a quantitative measure of tissue [^18^F][FDG uptake normalized for injected tracer dose and body weight, while the x-axis indicates the anatomical regions or tissues analyzed. White circles represent individual control participants; a blue circle represents Participant 2, the individual with KLEFS1 without a history of regression; and orange symbols represent individuals with KLEFS1 with a history of regression: Participant 1, circle; Participant 3, square; Participant 4, triangle; and Participant 5, diamond. Solid gray bars summarize the control group mean, with error bars indicating the standard error of the mean (SEM). [^18^F]FDG uptake was quantified as the mean SUV within spherical regions of interest (ROIs) placed in each tissue. Bilateral brain and muscle measurements were averaged before analysis. For Participant 4, who underwent two brain [^18^F]FDG-PET/CT scans approximately one year apart, values from both scans were averaged to generate a single value for analysis.

### Peripheral [^18^F]FDG metabolic profiles in KLEFS1

In addition to brain analyses, [^18^F]FDG uptake was assessed in liver, spleen, blood pool (left ventricle of the heart and ascending aorta), bone marrow (femoral corpus, L3 and L4 vertebral bodies), muscle (psoas major, deltoid, gluteus maximus), and adipose tissue (subcutaneous, visceral, and brown adipose tissue) for Participants 1-3 **(Table S2; Fig. 1C)**. Participant 2, similar to their brain metabolic findings, again exhibited generalized hypermetabolism, with increased [^18^F]FDG uptake in the liver, spleen, left ventricle, and L4 vertebral body **(Fig. 1C)**. Interestingly, elevated splenic uptake was observed in all participants. Additionally, Participant 1 showed increased uptake in the left ventricle, and Participant with borderline increased uptake in the L4 vertebral body. Muscle uptake tended to be lower than in controls, with Participants 1 and 3 showing reduced [^18^F]FDG accumulation in the deltoid, and Participant 3 exhibiting reduced uptake in the psoas major. Adipose tissue metabolism was generally comparable to controls, except for Participant 3, who demonstrated decreased metabolic activity in visceral and brown adipose tissue. Taken together, brain and peripheral [^18^F]FDG patterns indicate that Participant 2 shows a clear hypermetabolic profile, whereas the remaining participants show a tendency toward a hypometabolic state predominantly in the brain but also in select peripheral tissues. Although based on limited numbers, this contrast suggests distinct metabolic patterns between individuals with and without clinical regression, with hypermetabolism evident in the non-regression case and hypometabolism observed in those who experienced regression.

### KLEFS1 adipose tissue quantification

Body composition was analyzed on low-dose CT scans using a body composition algorithm which automatically identifies the axial cross-section at the midpoint of the third lumbar vertebra (L3). Cross-sectional areas of smooth muscle (SM), subcutaneous adipose tissue (SAT), visceral adipose tissue (VAT), and intermuscular adipose tissue (IMAT) were quantified for Participants 1, 2 and 3 **(Table S3; Fig. 2)**. Ages of Participants 1 and 3 were higher than those of the controls, and BMI was considerably elevated in Participants 1 and 2 **(Table S3)**. Smooth muscle and intramuscular adipose tissue were comparable to controls in all three individuals **(Fig. 2B,E)**. In contrast, Participants 2 and 3 showed increased subcutaneous adipose tissue and elevated SAT and SAT/ BMI ratios **(Fig. 2C,H)**. This was most notable in Participant 3, who had markedly increased SAT of 312 cm² (controls: 154 cm²) and a SAT/BMI ratio of 12.2 (controls: 6.3), despite having a BMI of 25.6 that was similar to the control average. Participant 2 additionally demonstrated increased visceral adipose tissue and an elevated VAT to BMI ratio **(Fig. 2G)**, consistent with their clinical obesity.

**Figure 2.**
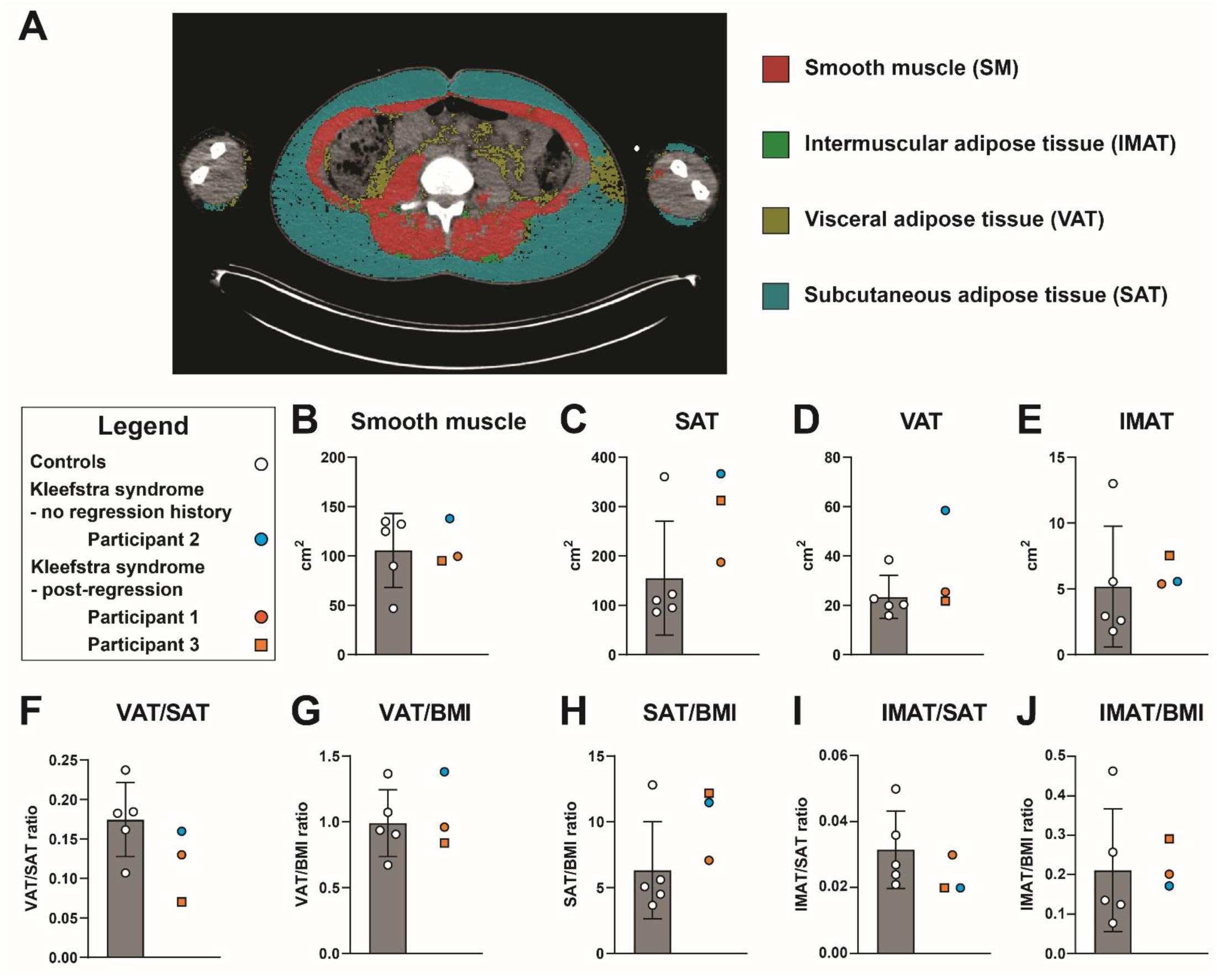
Quantification of abdominal body composition at the L3 vertebral level in individuals with KLEFS1 and controls. (A) Representative axial abdominal CT image (Participant 1) at the midpoint of the third lumbar vertebra (L3) with automated tissue segmentation generated by a body composition algorithm. Cross-sectional areas of smooth muscle (SM; red), subcutaneous adipose tissue (SAT; blue), visceral adipose tissue (VAT; olive), and intermuscular adipose tissue (IMAT; green) were quantified. (B–E) Cross-sectional areas (cm²) of SM, SAT, VAT, and IMAT. (F–J) Body composition ratios, including VAT/SAT, VAT/BMI, SAT/BMI, IMAT/SAT, and IMAT/BMI. The y-axis shows tissue area (cm²) or the indicated ratio, while the x-axis represents the study groups. Gray bars indicate the control group mean, with error bars representing the standard deviation (SD). White circles represent individual control participants, blue circles represent the individual with KLEFS1 without a history of regression (Participant 2), and orange symbols represent individuals with KLEFS1 with a history of regression (Participant 1, circle; Participant 3, square).

### KLEFS1 fly model displays baseline increased whole-body metabolism and neuronal ATP levels

To determine whether the metabolic alterations observed in KLEFS1 reflect conserved, mechanistically driven effects of *EHMT1* loss, we next examined baseline metabolic function in a well-established *Drosophila* model^32^. Previously, we characterized the *Drosophila* ortholog of *EHMT1*, *G9a*, as a critical regulator of metabolism in the fly brain and in other metabolically active peripheral tissues, including insulin-producing cells and the fat body^23,29^. We therefore asked whether baseline whole-body metabolic rate is also altered in the KLEFS1 fly model. Measuring whole-body CO_2_ production offers a reliable indicator of metabolic rate and can be readily quantified in freely moving flies^33^. Through respirometry, we analyzed CO_2_ production in both *G9a* mutants and their isogenic controls. Notably, *G9a* hemizygous mutant males exhibited elevated CO_2_ compared to the controls, thereby indicating an increase in metabolic rate due to the loss of *G9a* **(Fig. 3A)**.

**Figure 3.**
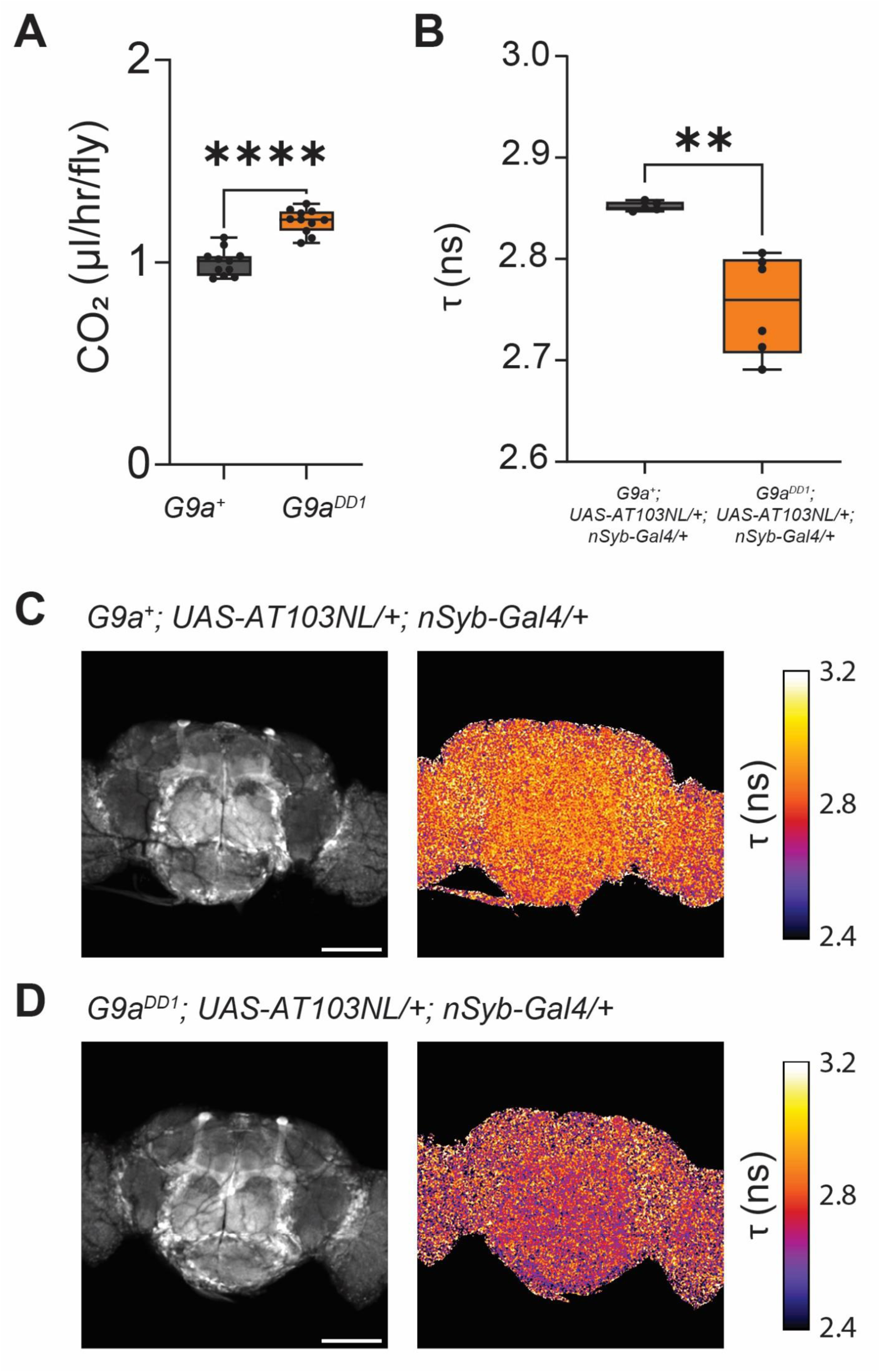
*G9a* loss leads to increased metabolic rate and ATP levels. **(A)** *G9a^DD1^* hemizygous mutant male flies (orange, n = 11) have increased CO2 production (P < 0.0001) compared to isogenic controls (*G9a^+^*, grey, n = 11). Each n represents the data from a single respirometer containing 5 animals of the same genotype. Box-and-whisker plots represent median, 25th–75th percentiles (box), and minimum-to-maximum values (whiskers). Statistical analysis was conducted using a two-tailed unpaired student t-test. (B) Fluorescence lifetime (τ) of the genetically encoded ATP sensor AT1.03NL was measured in pan-neuronal expression (*nSyb-Gal4*) brains from *G9a* hemizygous mutant males (*G9a^DD1^; UAS-AT1.03NL/+; nSyb-Gal4/+,* orange, n = 6) and isogenic controls (*G9a^+^; UAS-AT1.03NL/+; nSyb-Gal4/+*, grey, n = 5). Lower fluorescence lifetime values indicate higher intracellular ATP levels in *G9a* mutants (**p=0.0018). Box-and-whisker plots represent median, 25th–75th percentiles (box), and minimum-to-maximum values (whiskers). Statistical significance was assessed using a two-tailed unpaired Student’s t-test. (C) Representative central brain images of control flies (*G9a^+^; UAS-AT1.03NL/+; nSyb-Gal4/+*) and (D) *G9a* hemizygous mutants (*G9a^DD1^; UAS-AT1.03NL/+; nSyb-Gal4/+*) show fluorescent intensity (left panel) and the corresponding fluorescence lifetime map (right panel). Scale bars represent 100 µm.

Next, we assessed whether elevated whole-body metabolic rate in *G9a* mutants is accompanied by altered energy availability specifically within the brain. To measure ATP levels, male *G9a* mutants and isogenic controls expressed the genetically encoded ATP fluorescence resonance energy transfer (FRET) sensor AT1.03NL pan-neuronally under *nSyb-Gal4* control. AT1.03NL is a variant of the ATeam sensor that is optimized for use in *Drosophila*^34^. Brains were rapidly dissected in haemolymph-like buffer, which provides physiological ionic conditions suitable for *ex vivo* neuronal imaging and imaged by fluorescence-lifetime microscopy (FLIM). ATP binding to the sensor induces a conformational change that enhances FRET efficiency and correspondingly shortens donor fluorescence lifetime; thus, reduced lifetime values reflect higher intracellular ATP levels. Consistent with this relationship, *G9a* mutant brains exhibited significantly decreased fluorescent lifetimes relative to controls, indicating an overall increase in neuronal ATP levels **(Fig. 3B)**. Taken together, these findings demonstrate that loss of *G9a* leads to increased whole-organism metabolic rate and elevated brain ATP levels, paralleling the hypermetabolic phenotype observed in Participant 2 of the KLEFS1 cohort, who did not exhibit a regressive episode.

### Establishing a stress induced KLEFS1 regression paradigm

*G9a* mutant flies exhibit profound sensitivity to OS, characterized by rapid mortality linked to energy wasting and depletion of available energy stores^23^. Given our hypothesis that metabolic collapse under stress underlies regression in KLEFS1, we sought to establish an experimental paradigm that models a transient stress event followed by a recovery phase, allowing downstream phenotypes to be assessed independently of acute lethality. To this end, we used paraquat, a well-established inducer of OS^23,35^, and compared continuous exposure conditions with a transient exposure paradigm. Male *G9a* mutant and control flies were collected after eclosion, group housed for three days and then exposed to paraquat-containing food for 24 hours before transfer back to standard medium, hereafter referred to as paraquat-recovery treatment (PQ-R), while parallel cohorts were maintained under continuous paraquat exposure **(Fig. 4A)**.

**Figure 4.**
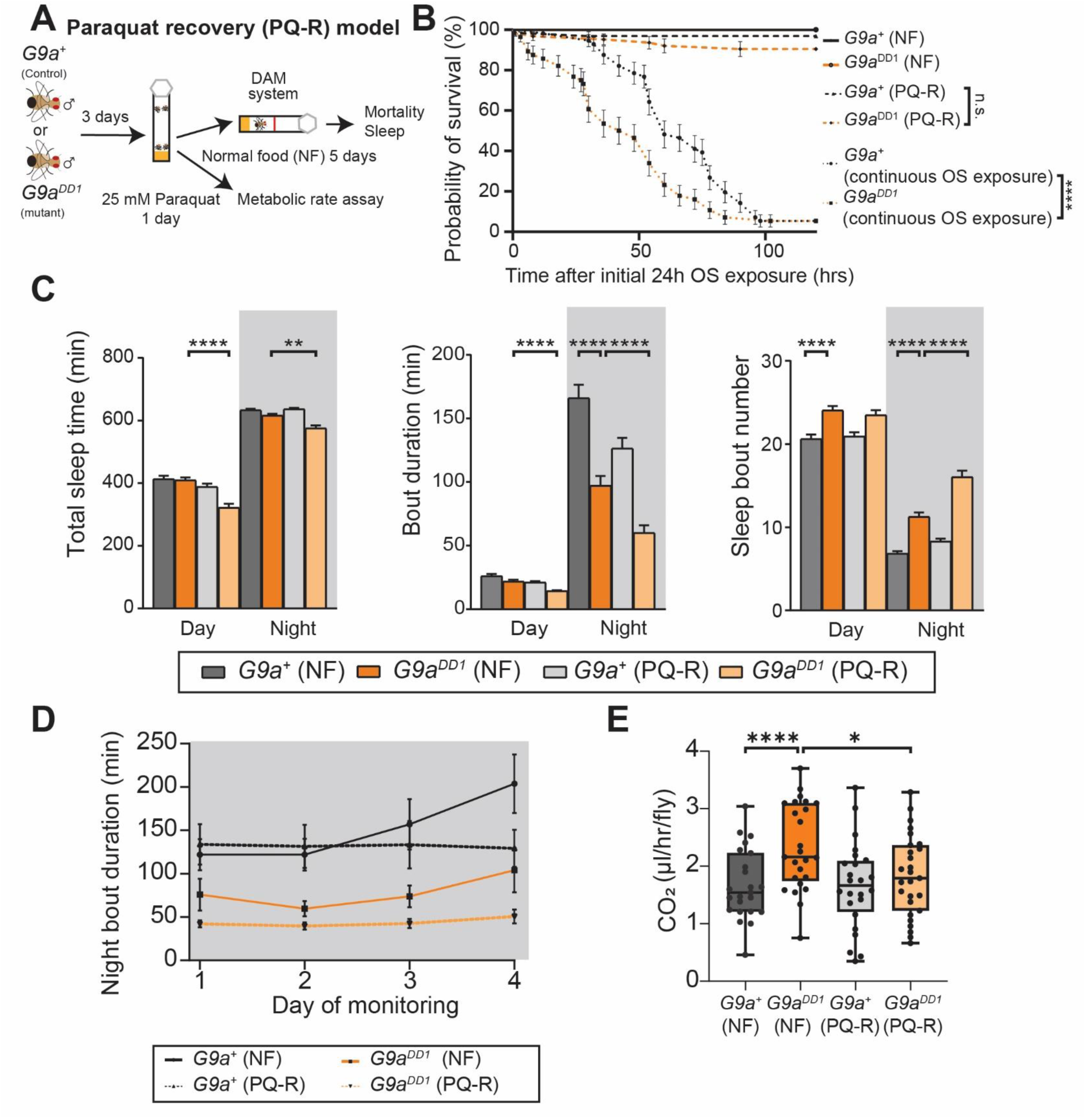
Transient oxidative stress induces lasting sleep disruption and metabolic shifts in *G9a* mutants. (A) Schematic of the paraquat-recovery (PQ-R) paradigm. Male *G9a* mutants (*G9a^DD1^*) and isogenic controls (*G9a^+^*) were group-housed for 3 days post-eclosion and exposed to paraquat-supplemented food (25 mM) for 24h before transfer back to standard medium (NF). (B) Survival curves under continuous paraquat exposure show significantly reduced survival in *G9a* mutants (n = 56) compared with controls (n = 56; p < 0.0001, Gehan–Breslow–Wilcoxon test). Under the 24 h PQ-R paradigm, survival of *G9a* mutants (n = 63) and controls (n = 64) did not differ from each other or from untreated cohorts, all of which remained fully viable over the monitoring period (mutant and controls, n=64). (C) Quantification of total sleep time, sleep-bout duration, and sleep-bout number during the day and night under non-food (NF) and PQ-R conditions. Under NF conditions, *G9a* mutants (dark orange, n = 127) show fragmented nighttime sleep, with reduced bout duration and increased bout number, compared to control flies (dark grey, n=125), while total sleep is maintained. Following PQ-R treatment, G9a mutants (light orange, n = 85) show further reductions in both daytime and nighttime sleep-bout duration (****p < 0.0001), accompanied by increased nighttime bout number (****p < 0.0001), resulting in significantly disrupted total sleep across both day and night (****p < 0.0001, **p = 0.0051). Control flies (light grey, n = 113) show a non-significant trend toward sleep fragmentation. Data are presented as box plots with mean ± SEM. Statistical analyses were conducted using Kruskal–Wallis tests followed by Dunn’s multiple comparisons test. Bonferroni correction was then applied across genotype-specific datasets and corrected two-sided significance thresholds were used to determine statistical significance. (D) Nighttime sleep-bout duration across four days of monitoring following PQ-R treatment. *G9a* mutants (n=35 (PQ-R), n=50 (NF)) exhibit a persistent reduction in nighttime bout duration that remains stable over the monitoring period, in contrast to controls (n=52 (PQ-R), n=60 (NF)). (E) Whole-body metabolic rate (CO₂ production) following PQ-R treatment. *G9a* mutants (n = 22 (PQ-R), n=21 (NF) respirometers; 5 flies each) display a significant reduction in CO₂ output (*p = 0.0224), shifting from a hypermetabolic baseline (****p < 0.0001) toward control levels, whereas control flies (n = 20 (PQ-R), n=19 (NF)) show no significant change. Box-and-whisker plots represent median, 25th–75th percentiles (box), and minimum-to-maximum values (whiskers). Normality was assessed prior to analysis. Normally distributed datasets were analyzed using two-way ANOVA followed by Tukey’s multiple comparisons test.

As expected, under continuous exposure to paraquat, *G9a* mutants exhibited significantly reduced survival compared to controls, with 50% mortality occurring at approximately 74 hours after OS onset **(Fig. 4B)**. In contrast, following 24-hour PQ-R treatment, survival of *G9a* mutants and controls was not significantly different from each other or from unexposed flies, with all groups exhibiting high survival rates throughout the monitored period. These results demonstrate that a 24-hour physiological challenge of paraquat-induced OS in *G9a* mutants does not result in lasting reductions in survival, enabling subsequent analysis of post-stress behavioural and metabolic phenotypes relevant to regression in KLEFS1.

### KLEFS1 regression model displays exacerbated sleep disruption and altered metabolic rate

*G9a* mutant flies display sleep disturbances that parallel those reported in KLEFS1 and this has been linked to *G9a*’s role in maintaining reactive oxygen species (ROS) homeostasis within metabolically active tissues^29^. Nighttime sleep plays a critical role in clearing ROS that accumulate during the day^36,37^, and disrupted nighttime sleep is a major concern reported by caregivers of individuals with KLEFS1^11^. With this in mind, we examined whether the transient OS challenge in our PQ-R paradigm would exacerbate these sleep defects, with particular focus on nighttime sleep. Under standard conditions, *G9a* mutants exhibited fragmented nighttime sleep, characterized by reduced sleep bout duration and a compensatory increase in sleep bout number, resulting in no significant change in total sleep time, however, indicative of overall reduced sleep integrity **(Fig. 4C)**. Following PQ-R treatment, *G9a* mutants showed a further reduction in both daytime and nighttime sleep bout duration, with partial compensation by increased nighttime bout number but not during the day. This resulted in disrupted total sleep across both day and night, reflecting a further deterioration of sleep architecture. While control flies showed a trend toward increased sleep fragmentation following paraquat exposure, these changes were not statistically significant. Notably, the reduction in nighttime sleep bout duration remained stable in *G9a* mutants across the monitoring period, indicating a persistent post-stress effect that lasts at least for four days **(Fig. 4D)**.

From our clinical cohort, individuals who had experienced developmental regression displayed a distinct metabolic profile characterized by hypometabolism. This prompted us to examine whether whole-body metabolism was similarly affected in our KLEFS1 regression model. Following PQ-R treatment, we found that metabolic rate in *G9a* mutants was decreased significantly, shifting from a hypermetabolic state back toward the control baseline, whereas controls showed no significant change **(Fig. 4E)**. This stress-induced metabolic shift echoes the altered metabolic state observed in individuals who had experienced regression in the KLEFS1 cohort. While not directly equivalent to the hypometabolic pattern observed in KLEFS1, this shift, together with the exacerbated and persistent sleep disruption, indicates that transient OS induces lasting changes in behavioural and metabolic phenotypes in the KLEFS1 regression model.

### Adult high-sugar feeding mitigates OS-induced sleep and metabolic disruption in the KLEFS1 regression model

High-sugar diet (HSD) has previously been shown to enhance OS resistance in the *Drosophila* KLEFS1 model by increasing accessible energy stores, thereby improving survival during metabolic challenge^23^. Given that transient OS exposure exacerbated nighttime sleep fragmentation in our KLEFS1 regression model, we asked whether increasing energy availability through dietary sugar supplementation could also buffer *G9a* mutants from these OS-induced sleep defects. To test this, we implemented three HSD regimens - continuous exposure, developmental-only exposure, and adult-only exposure - and assessed their impact on nighttime sleep architecture, with a specific focus on sleep bout duration, under baseline and PQ-R conditions.

Across all dietary assays, *G9a* mutants exhibited disrupted nighttime sleep at baseline compared to controls, consistent with our earlier findings, and PQ-R exposure further exacerbated this fragmentation under standard-diet feeding. Continuous HSD feeding increased nighttime sleep bout duration in both controls and *G9a* mutants by approximately 50% under baseline conditions, potentially indicating enhanced sleep integrity when energy availability is chronically elevated **(Fig. 5A)**. However, despite this, continuous HSD did not prevent the OS-induced decrease in nighttime sleep bout duration following PQ-R treatment. Similarly, developmental-only HSD failed to mitigate sleep disruption, as *G9a* mutants still showed a significant reduction in bout duration following PQ-R exposure **(Fig. 5B)**.

**Figure 5.**
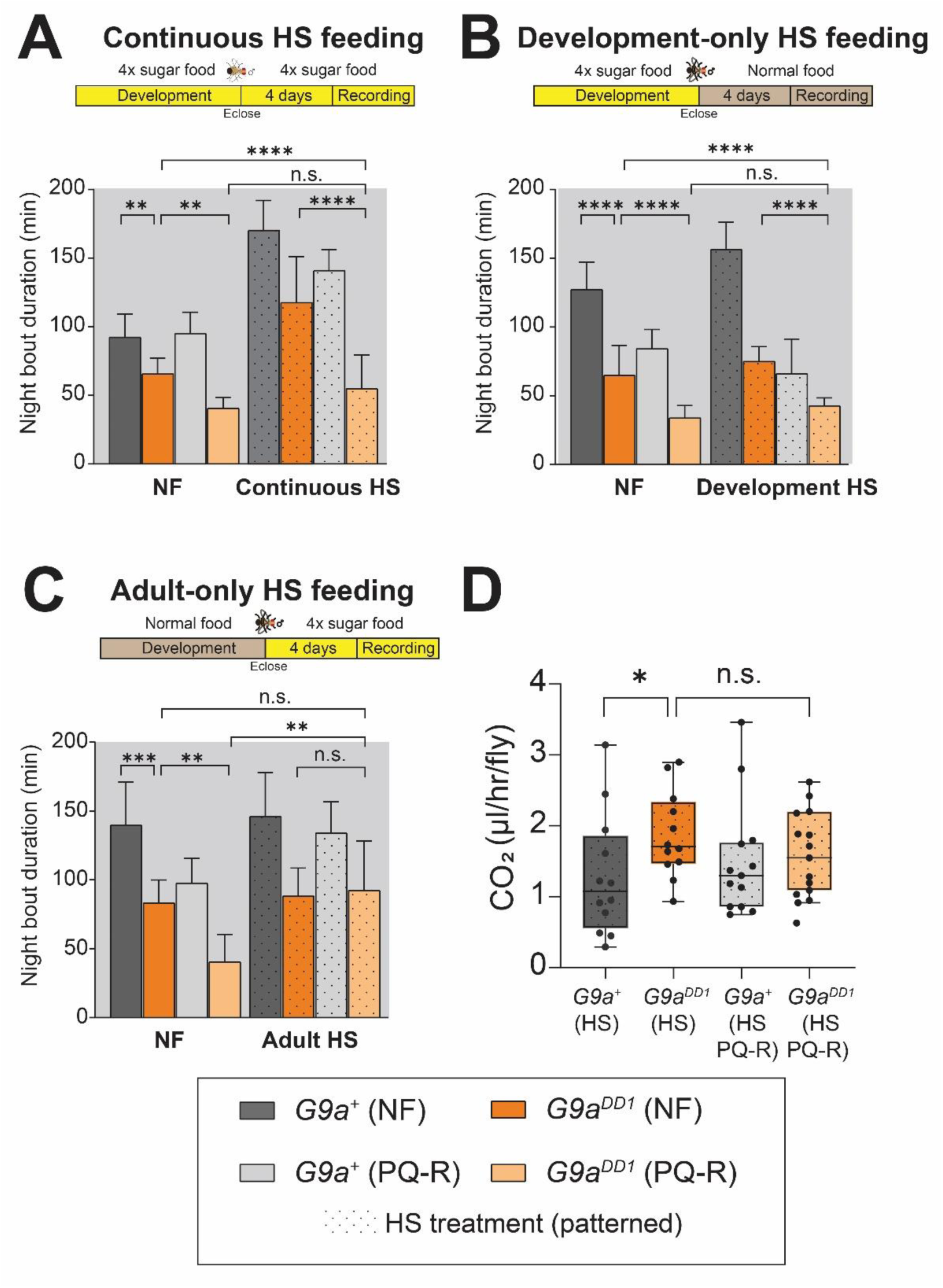
Adult high-sugar feeding mitigates OS-induced sleep and metabolic disruption in the KLEFS1 regression model. (A) Continuous high-sugar (HS) feeding. Nighttime sleep-bout duration (grey background) in *G9a* mutants (*G9a^DD1^*) and isogenic controls (*G9a^+^*) under normal food (NF) and continuous HS conditions, with and without paraquat-recovery (PQ-R). HS feeding increases baseline sleep-bout duration in both genotypes (*G9a^DD1^*: dark orange, n = 88 (NF), dark orange with dots, n = 92 (HS); *G9a^+^*: dark grey, n = 84 (NF), dark grey with dots n = 67 (HS)). PQ-R significantly reduces sleep-bout duration relative to HS baselines in both genotypes (*G9a^DD1^*: light orange, n = 67 (PQ-R), light orange with dots n = 34 (HS-PQ-R); *G9a^+^*: light grey, n = 88 (PQ-R), light grey with dots, n = 63 (HS-PQ-R)). Data are presented as box plots with mean ± SEM. (Kruskal– Wallis with Dunn’s multiple comparisons; *p < 0.05, **p < 0.01, ***p < 0.001, ****p < 0.0001). (B) Developmental-only HS feeding. Sleep-bout duration under NF and developmental HS conditions followed by PQ-R. Developmental HS does not prevent PQ-R-induced sleep fragmentation in G9a mutants (*G9a^DD1^*: n = 55 (NF), n = 54 (Dev-HS), n = 39 (PQ-R), n=41 (Dev-HS-PQ-R); *G9a^+^*: n = 56 (NF), n = 47 (Dev-HS), n = 54 (PQ-R), n=36 (Dev-HS-PQ-R). Statistics as in (A). (C) Adult-only HS feeding. Sleep-bout duration under NF and adult HS conditions with PQ-R exposure. Unlike other regimens, adult HS preserves sleep-bout duration in G9a mutants following PQ-R, preventing OS-induced fragmentation (*G9a^DD1^*: n = 55 (NF), n = 57 (Adult-HS), n = 36 (PQ-R), n=45 (Adult-HS-PQ-R); *G9a^+^*:n = 60 (NF), n = 62 (Adult-HS), n = 49 (PQ-R), n=58 (Adult-HS-PQ-R). Statistics as in (A). (D) Whole-body metabolic rate following PQ-R. CO₂ production in *G9a^DD1^* and controls under NF and adult HS conditions. *G9a* mutants remain elevated at baseline (*p = 0.0137), including under HS (*G9a^DD1^*: dark orange, n =12, *G9a^+^*: dark grey, n = 12, respirometers; ∼5 flies per respirometer). Adult HS prevents PQ-R-induced metabolic decline, maintaining levels comparable to unexposed mutants (*G9a^DD1^*: light orange, n =14, *G9a^+^*: light grey, n = 13, HS-PQ-R). Data are shown as box plots (median, 25th–75th percentiles; whiskers min-max). Metabolic data were analyzed after normality testing using two-way ANOVA followed by Tukey’s multiple comparisons test .

By contrast, adult-only HSD produced a strikingly different outcome. Here, *G9a* mutants did not exhibit the characteristic PQ-R-induced decline in nighttime sleep bout duration, indicating that adult-restricted high-sugar feeding protected nighttime sleep from OS-induced deterioration **(Fig. 5C)**. Notably, unlike continuous HSD, adult-only HSD did not increase nighttime sleep at baseline in *G9a* mutants and controls. This distinction suggests that the elevated baseline nighttime sleep seen in *G9a* mutants and control under continuous HSD may reflect broader physiological effects of chronic high-sugar exposure, rather than true improvements in sleep integrity, whereas adult-only HSD uniquely confers protection during acute metabolic challenge.

Given this protective effect on sleep, we then examined whether adult-only HSD similarly buffered whole-body metabolic rate in the context of OS exposure. *G9a* mutants remained hypermetabolic under baseline conditions, even when maintained on HSD feeding **(Fig. 5D)**. However, following PQ-R treatment, adult-only HSD prevented the significant reduction in metabolic rate observed in standard-fed *G9a* mutants, resulting in no significant difference from unexposed mutants.

Together, these findings indicate that adult-restricted high-sugar feeding confers metabolic resilience during transient OS exposure, preserving both metabolic rate and nighttime sleep architecture. This supports the model that regression-related phenotypes in KLEFS1 arise from an inability to access or mobilize energy upon metabolic challenge, and that targeted metabolic support in adulthood can selectively buffer against the behavioural consequences of transient OS exposure.

## Discussion

Developmental regression represents one of the most severe and least understood complications of KLEFS1 and other neurodevelopmental disorders. In this study, we identified alterations in cerebral and peripheral glucose metabolism in individuals with KLEFS1 and used a *Drosophila* G9a loss-of-function model to explore whether metabolic stress may contribute to regression-related phenotypes. [^18^F]FDG-PET/CT revealed frontal brain hypometabolism in four of five individuals, including all individuals with a reported history of regression, whereas one individual without regression showed widespread hypermetabolism. Although these findings are based on a small exploratory cohort and do not establish a relationship between cerebral hypometabolism and regression, the observed pattern raises the possibility that altered cerebral glucose metabolism may be associated with regression susceptibility. Alongside these central findings, all three individuals who underwent whole-body imaging showed increased glucose uptake in peripheral immune-related tissues, including the spleen and, in some cases, the blood pool and vertebral bone marrow. These observations provide preliminary evidence of altered glucose metabolism across central and peripheral tissues in KLEFS1 and raise the possibility of metabolic and immune-related alterations in the disorder.

In the *Drosophila* model, *G9a* mutants displayed increased baseline metabolic rate and neuronal ATP levels, together with fragmented nighttime sleep. Oxidative stress further exacerbated sleep fragmentation and reduced metabolic output in the mutants, providing experimental evidence that *G9a* deficiency may increase susceptibility to metabolic disruption under environmental stress. Notably, high-sugar feeding during adulthood mitigated stress-induced sleep deterioration and helped maintain metabolic output. These findings suggest that metabolic state may influence the response to environmental stress in *G9a* mutants and provide a potential framework for interpreting the metabolic abnormalities observed in individuals with KLEFS1. Together, these complementary human and *Drosophila* findings support the hypothesis that altered metabolic regulation may contribute to vulnerability to regression-related phenotypes in KLEFS1, while further studies will be required to establish whether and how these metabolic alterations relate to regression in affected individuals.

### KLEFS1 and dysregulated immunometabolic function

Converging evidence from neurodevelopmental and neurodegenerative disorders underscores the sensitivity of cerebral glucose metabolism to synaptic, mitochondrial, and inflammatory dysfunction. Brain glucose hypometabolism, as observed in the four individuals with a history of regression, is a feature shared with Alzheimer’s disease and Parkinson’s disease, both of which involve impaired glucose utilization linked to mitochondrial dysfunction and insulin resistance^38^. Hypometabolism in schizophrenia, particularly in frontal regions and basal ganglia, has similarly been associated with synaptic dysfunction and decreased energetic demand^39^. These comparisons support a model in which frontal hypometabolism in KLEFS1 may reflect decreased synaptic activity or impaired metabolic capacity, although the underlying mechanisms remain to be determined.

Conversely, the single individual with widespread hypermetabolism mirrors patterns observed in conditions such as Rett syndrome, where elevated glutamate– glutamine cycling increases astrocytic glucose uptake^40^, or Down syndrome, where hypermetabolism has been interpreted as compensatory neuronal activity in underdeveloped networks^41^. Hypermetabolism also appears in certain metabolic disorders and inflammatory or immune-mediated brain diseases^42^. These precedents raise the possibility that the hypermetabolic profile observed in the non-regressive individual reflects a distinct metabolic state within KLEFS1, one that may characterize individuals who have not undergone regression, although additional cases will be needed to determine whether this pattern holds more broadly. Taken together, these clinical patterns suggest that metabolic alterations in KLEFS1 may not follow a uniform direction but instead reflect altered energetic balance. The direction of metabolic change likely depends on state, timing, and the history of environmental or physiological stress, raising the possibility that hyper- and hypometabolic profiles may represent different positions along a shared vulnerability trajectory rather than categorically distinct subtypes.

Across all individuals, increased [^18^F]FDG uptake in the spleen is consistent with increased immune-related metabolic activity. Increased spleen uptake can reflect immune activation, elevated hematopoietic demand, or abnormalities in glucose metabolism, but none of the individuals had autoimmune disease, anemia, or diabetes mellitus, and only one exhibited insulin resistance. Increased blood pool uptake, observed particularly in the right ventricle, may similarly reflect systemic metabolic activation of immune cells. These findings align with evidence that KLEFS1 involves overactivation of the inflammasome pathway and increased inflammatory status^43^. Together, these findings raise the possibility that altered metabolic activity extends across both neural and immune-related tissues in KLEFS1.

In our *Drosophila* KLEFS1 model, we found that *G9a* mutants exhibit disrupted systemic metabolic regulation consistent with the metabolic alterations observed in individuals with KLEFS1. At baseline, *G9a* mutants display an increased whole organism metabolic rate and elevated neuronal ATP levels. The concurrent increase in ATP levels despite elevated metabolic demand suggests increased energetic turnover, in which ATP production is over-upregulated to match higher consumption. As our measurements capture systemic metabolic output and overall neuronal ATP levels, future work is required to resolve tissue-, organelle-, and pathway-level contributions. It is possible that increased oxidative phosphorylation, required to sustain elevated ATP levels, contributes to increased reactive oxygen species production and potentially reflect underlying mitochondrial stress^44^. Consistent with this, these baseline metabolic differences are accompanied by altered redox homeostasis in the KLEFS1 fly model, including elevated methionine sulfoxide levels^29^, indicating an underlying imbalance in reactive oxygen species handling. This pre-existing metabolic fragility may partially contribute to the increased sensitivity of *G9a* mutants to oxidative stress^23^, which was used here to model regression like conditions. Importantly, metabolic systems often undergo stress-induced reprogramming, including a shift from oxidative phosphorylation toward aerobic glycolysis (the Warburg effect), which can support rapid energetic and biosynthetic demands during immune activation, but if prolonged, this reprogramming drains resources and results in significant energy wasting^45^. Following oxidative stress exposure, we found that mutants show a downward shift in whole organism metabolic rate, reflecting a loss of metabolic stability under environmental challenge and paralleling the regression associated metabolic changes observed in KLEFS1. Mechanistically, this metabolic vulnerability to energy wasting aligns with the role of G9a as an epigenetic regulator that limits excessive transcriptional activation in response to environmental stress, thereby buffering energetically costly gene expression programs^23,24^. In this context, impaired regulatory control may promote maladaptive stress responses that couple immune activation and metabolic demand, providing a potential link between metabolic instability and stress-induced changes in sleep and metabolic output. Taken together, these findings support the hypothesis that altered stress-sensitive metabolic regulation may contribute to vulnerability to environmental challenges in KLEFS1.

### KLEFS1 regression at the intersection of sleep and ROS-linked metabolism

Sleep disturbances are highly prevalent in neurodevelopmental disorders and are a core feature of KLEFS1. Our prior work shows that *G9a* mutants exhibit fragmented sleep arising from metabolic dysfunction in insulin producing cells and fat body, accompanied by elevated ROS related metabolites^29^. This positions EHMT1/G9a as a regulator of sleep integrity through control of ROS homeostasis. ROS generated during oxidative phosphorylation accumulate during wake and are normally neutralized during sleep^36^, while disruption of this balance promotes both sleep instability and oxidative stress accumulation. Multiple studies have shown that prolonged sleep deprivation leads to an antioxidant response in the brain^46–48^ and ROS accumulation in the gut^49^. Moreover, increasing ROS with a mild dose of paraquat results in sleep fragmentation^50^, a pattern our PQ-R control flies also trended toward, whereas fragmenting sleep leads to an increase of ROS^51^. *Drosophila* short sleeping mutants are sensitive to oxidative stress, while increasing sleep in wild-type flies promotes oxidative stress resistance^52^.

In this study, we leveraged this framework to model regression by applying oxidative stress to exacerbate baseline sleep fragmentation. We focused on nighttime sleep bout duration as a measure of sleep architecture, reflecting the importance of consolidated sleep episodes rather than total sleep time for neural stability and cognitive function^53^. Here we found that oxidative stress further disrupted sleep architecture in *G9a* mutants and induced a correlated reduction in metabolic rate, consistent with a failure to maintain metabolic resilience under conditions of increased oxidative burden. To test whether extra energy availability could buffer oxidative stress induced sleep disruption, we provided a high sugar diet immediately after eclosion through adulthood. Excitingly, adult restricted high sugar feeding prevented the paraquat induced exacerbation of nighttime sleep fragmentation in *G9a* mutants and the associated reduction in metabolic rate. Whether this protection also extends to other regression linked phenotypes such as memory deficits already present at baseline in KLEFS1 models^32^ remains an open question.

Interestingly, a continuous high-sugar exposure did not confer protection to prevent OS-induced sleep deterioration. Chronic high sugar diets have been shown to disrupt sleep architecture and increase fragmentation^54,55^, and in our paradigm increased nighttime sleep bouts, highlighting a general remodeling of sleep under increased energy availability. These differences we see between adult-only and continuous high sugar diets may reflect how short term versus chronic high sugar exposure engages the gut-brain axis, with long term feeding inducing gut inflammation that disrupts sleep^54^, whereas short term supplementation can suppress age-related sleep fragmentation^55^. Together, these findings raise the possibility that targeted metabolic support could influence stress-induced phenotypes relevant to regression in KLEFS1.

Given the emerging role of gut-brain metabolic signaling in shaping sleep and oxidative stress responses^54^, additional cellular regulators of ROS buffering warrant consideration in the context of KLEFS1. While our prior work identified fat body and insulin producing cells as critical nodes for *G9a* dependent sleep regulation^29^, the growing evidence for glial contributions to metabolic support and redox homeostasis, together with the identification of neuroinflammatory astrocyte dysfunction in EHMT1-deficient cell models as a therapeutic target of olanzapine during developmental regression^17^, makes them a compelling additional target for understanding ROS buffering and energy balance relevant to regression. During sleep, glia clear and metabolize peroxidated lipids that accumulate during wakefulness^37^, a process dependent on coordinated metabolic support. This is further reinforced by cross tissue energy regulation pathways linking glia, fat body, and the gut, where Gart, a trifunctional enzyme involved in purine biosynthesis and metabolic regulation, coordinates feeding rhythms and energy storage via lipid and glycogen regulatory programs^56^. Importantly, key components of these pathways including *Bmm*, *Lsd-2*, *Ugp*, and *Gart* itself are dysregulated in *G9a* mutants under stress conditions^23^. Given that *Gart* is also under circadian control via the CLK/CYC heterodimer, it will also be of interest to determine whether G9a-dependent chromatin regulation intersects with rhythmic metabolic control. Altogether, these findings motivate a model in which disrupted sleep architecture, ROS imbalance, and altered metabolic coordination across neural and peripheral tissues may interact to influence vulnerability to regression in KLEFS1.

### Limitations

Studying the clinical aspects of individuals with severe rare diseases comprising intellectual disability and psychiatric disorders such as described in this study, is extremely challenging. Both in terms of the numbers that can be included and in terms of systematic approaches. Though we describe a limited number of five cases, the observations in this cohort warrant clinical follow up in further patients.

Numerous factors influence [^18^F]FDG-PET/CT results, including BMI, fasting state, glucose levels, insulin, physical activity, and timing of imaging^57^. For some participants, glucose levels at the time of the scan were unavailable, though available measurements were within normal limits. Medication and anesthetic exposure represent additional potential confounders, as antipsychotics and anesthetics such as propofol can reduce brain glucose metabolism^31,39,58,59^. Two participants received olanzapine for psychosis which accompanied regression, which may therefore have contributed to reduced cerebral glucose uptake. However, individuals who had experienced regression not receiving these medications also exhibited hypometabolism, suggesting that medication effects are unlikely to fully account for the observed findings. The potential peripheral effects of these treatments remain unknown. Finally, the lack of full-brain imaging in controls may affect comparisons for some regions close to the marginal edge.

On the *Drosophila* side, oxidative stress is a simplification of the environmental stresses that may precipitate regression in humans, and sleep architecture cannot be directly equated across species. Nevertheless, the observation of metabolic alterations in both the clinical cohort and *Drosophila* model provides complementary evidence and supports further investigation of metabolic regulation as a potential contributor to regression vulnerability.

## Methods

### Participant recruitment and study design

A cohort of five individuals with KLEFS1, and five healthy controls were analyzed in this study. Three individuals diagnosed with KLEFS1 received a 2-Deoxy-2-[^18^F]fluoro-D-glucose ([^18^F]FDG) PET scan, including low-dose CT for attenuation correction with a Siemens Biograph 40 mCT scanner (Siemens Healthcare) at the Radboudumc. After a low-carb diet and while under general anesthesia, a standard-of-care whole-body scan ranging from toes to the top of the skull was acquired. Dosing was performed according to EANM guidelines with intravenous injection of 2.1 MBq/kg (±10 %) after at least 4 hours fasten, drinking 500 mL water and serum glucose levels <8 mmol/L^60^. Scans from skull to mid-thigh were acquired at 60±5 minutes post-injection. Two other individuals received a standard-of-care brain [^18^F]FDG PET/CT scan at centers in France (Participants 4 and 5) with a Siemens Biograph 20 mCT scanner (Siemens Healthcare). Participant 4 received a scan at two different timepoints approximately one year apart.

The control scans were obtained as baseline during a previous study into BCG vaccination (METC #2015-2177^61^). Of this larger cohort of healthy subjects, the baseline (pre-vaccination) [^18^F]FDG PET/CT scans of five age and weight matched subjects were analyzed. Standard-of-care scan protocols were applied based on a scan range from the trochanter major to the base of the skull, and two controls received an extended scan range including all brain lobes.

All KLEFS1 diagnoses were molecularly confirmed (EHMT1 haploinsufficiency) before inclusion in this study. Information from medical files (age, sex, body mass index (BMI), medication use) was also collected for description and correction.

### Ethics

This study was approved under Statement #2024-17642 and is not subject to the Dutch Medical Research Involving Human Subjects Act (WMO). Retrospective data were extracted from the Radboudumc Biobank Genetics and Rare Disease (#2018-4985). Scans from France were shared with consent from participants’ legal representatives, and Dutch participants provided consent through biobank enrollment.

### [^18^F]FDG-PET/CT imaging and analyses

The primary outcome measure was the level of [^18^F]FDG uptake in spheroids of comparable size in tissues of interest in both groups. These spheroids represented spherical regions of interest (ROIs). Tracer uptake was quantified as mean Standardized Uptake Value (SUV) using IRW (Inveon Research Workplace 4.2, Siemens Molecular Imaging).

SUV was calculated as:

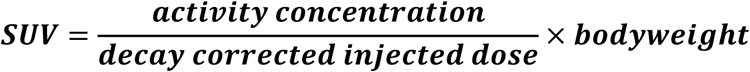

The level of tracer uptake was determined in the brain, various organs, muscles, blood pool, bone marrow, and adipose tissue. Brain and muscle measurements were obtained bilaterally, with left–right averages used for analysis.

Spheroids in the brain were placed in the frontal lobe, temporal lobe, basal ganglia, and cerebellum. Organ measurements included the liver and spleen. The blood pool was assessed in the left ventricle and ascending aorta. Bone marrow activity was measured in vertebral bodies L3 and L4 and in the femoral corpus. The mean of spheroids placed in the deltoid, psoas major, and gluteus maximus represented musculature. Adipose tissue measurements were obtained in brown adipose tissue (BAT), subcutaneous adipose tissue (SAT), visceral adipose tissue (VAT), and intramuscular adipose tissue (IMAT). Participants scanned in France received only a brain scan; therefore, peripheral tissue analysis could not be performed.

Using the body composition algorithm, accessed online through the grand-challenge.org platform (https://grand-challenge.org/algorithms/body-composition/), the quantity and ratio of SAT, VAT, IMAT and smooth musculature (SM) were analyzed based on a low-dose CT slice at the L3 level.

### Fly stocks and genetics

*Drosophila melanogaster* stocks were reared on a standard medium (yeast-cornmeal-agar-sugar), supplemented with the mold inhibitors methyl paraben and propanoic acid, at 25 °C in 70% humidity, and entrained with a 12h:12h light/dark cycle. The following *Drosophila* stocks were used in this study: *G9a* mutant flies (*G9a^DD1^*, previously described^32^) and their isogenic background controls, corresponding to the Vienna *Drosophila* Resource Center (VDRC) GD library (stock #60000). For ATP metabolic imaging experiments, we obtained a genetically encoded FRET-based ATP biosensor, AT1.03NL (DGRC #117011; genotype: w[*]; P{w[+mC]=UAS-AT1.03NL}1), which was expressed pan-neuronally using *nSyb-Gal4*. The UAS-AT1.03NL line was crossed to either *G9a* mutants or their genetic background controls to generate experimental and control progeny expressing the ATP sensor specifically in neurons.

### Paraquat exposure

Oxidative stress (OS) was induced using paraquat (methyl viologen dicholoride hydrate, 98 percent; Sigma 856177). A 1 M stock solution of paraquat was stored at −20°C and was diluted into freshly made food at 40°C to a final concentration of 25 mM^23^. Paraquat was added to both standard and high sugar diet media as required.

To model OS-induced regression in the KLEFS1 fly model, *G9a* mutant and isogenic control male flies were collected after eclosion and allowed to recover from CO_2_ anesthesia for three days. Flies were then transferred and exposed to paraquat-containing food for 24 hrs while incubated at 25°C and 70% humidity. Following exposure, flies were transferred to the appropriate assay conditions as described below.

### High sugar diet treatment

High sugar diet (HSD) treatment was implemented using food prepared from the standard recipe with a fourfold increase in sugar concentration (440g/L)^23^. Three exposure paradigms were used. In the developmental-only condition, *G9a* mutant and control flies were reared on HSD from egg to eclosion and then transferred to standard food immediately after eclosion. In the adult-only condition, flies were reared on standard food and transferred to HSD immediately after eclosion, with HSD exposure maintained throughout paraquat treatment and sleep monitoring. In the continuous-HSD condition, flies were reared on HSD throughout development and remained on HSD after eclosion and during the sleep monitoring phase. PQ-R summary analyses **(Fig. 4C)** were conducted using data collected from corresponding high-sugar dietary experiments.

### Sleep monitoring and survival analysis

Sleep and survival was monitored using the *Drosophila* Activity Monitor system (DAM2) system (Trikinetics, Waltham, MA, USA). In brief, four day old male flies were individually transferred without CO_2_ anesthesia into plastic tubes (65mm by 5mm) containing the appropriate food condition. For survival assays specifically, tubes contained either standard food or standard food supplemented with 25 mM paraquat, as required. Tubes were loaded into the DAM2 monitors and placed immediately in the incubator at 25 °C under a 12:12 light–dark cycle, initiating the recording period.

Flies were allowed to acclimate for at least 12 hours before sleep data collection. Activity was then recorded for four days. Movement was detected via infrared beam crossings, and data were processed using DAMFileScan110 (Trikinetics) and the Sleep and Circadian Analysis MATLAB Program (SCAMP)^62^. Sleep was defined as periods of five or more minutes of inactivity^63,64^, and sleep parameters represent averages across the four-day acquisition period.

For survival assays, monitoring continued for a total of 120 hours. Time of death was defined as the point at which activity reached zero.

### Metabolic rate (CO_2_) measurement

Metabolic rate was measured via respirometry as an index of CO_2_ production^65^. Custom-made respirometers were constructed by attaching a 50 µL capillary micropipette to a 1 mL pipette tip. After drying, each respirometer was filled with soda lime (#72073, Sigma-Aldrich) as a CO₂ absorbent, positioned between two cotton plugs to prevent direct contact with the flies and with either standard food or high sugar diet food as required. For each experimental condition, four day old male flies were used. Flies were cold anesthetized and groups of five individuals of the same genotype and treatment condition were transferred into each respirometer, followed by a 15-minute recovery period. Respirometers were sealed with modelling clay and mounted vertically on a polystyrene rack. One respirometer without flies was included as an atmospheric and temperature control. In total, ten respirometers were used per experiment, including one control and nine experimental units. Respirometers were placed inside a sealed chromatography chamber containing a bromophenol blue water-based solution (#B5525, Sigma-Aldrich) at the base. The chamber was closed with an airtight lid to minimize environmental variability. The setup was equilibrated in a behavioural room maintained at 25 °C and 70 percent humidity for 75 minutes prior to data collection (baseline). After equilibration, images were taken at baseline and again one hour later. CO₂ production was quantified by measuring the displacement of the liquid meniscus using Fiji software. The difference between baseline and final measurements was converted into volumetric CO₂ production and corrected for atmospheric and temperature fluctuations using the negative control respirometer, then normalized to fly number to yield metabolic rate expressed as µL CO₂ per hour per fly. Respirometers that failed to maintain an airtight seal (identified by absence of meniscus displacement in controls) were excluded from analysis.

### ATP imaging sample preparation and acquisition

Adult *G9a* mutant and control male flies expressing the genetically encoded ATP sensor AT1.03NL (DGRC #117011) under pan-neuronal *nSyb-Gal4* control were anaesthetized on ice and immediately transferred to ice-cold haemolymph buffer (HB)^66^. HB contained 130 mM NaCl, 5 mM KCl, 2 mM MgCl₂, 2 mM CaCl₂, 5 mM D-trehalose, 30 mM sucrose, and 5 mM HEPES-hemisodium salt. Brains were dissected in HB and transferred to the microscope within 20 minutes of dissection for fluorescence-lifetime imaging. Fluorescence lifetime measurements were acquired on a Leica SP8 microscope, using tuneable white-light laser excitation at 488 nm (80 MHz pulse frequency), and fluorescence was collected by time-correlated single-photon counting over 495–545 nm. For each brain, z-stacks were collected in six steps spanning 15 µm between the anterior and centre of the brain, then photon counts were summed across slices, and fluorescence lifetime (τ) was calculated using the FLIMJ plugin in ImageJ.

### Statistical analysis

All statistical analyses were performed using GraphPad Prism version 10.6.1. Sleep statistical analyses were performed as previously described^29,67^, with comparisons involving more than two genotypes using the Kruskal–Wallis test followed by Dunn’s multiple comparisons test. Where applicable, Bonferroni correction was applied to account for multiple testing across datasets per genotype, and corrected two-sided significance thresholds were used to determine statistical significance. Outliers were identified and removed using the ROUT method (Q = 1%). Survival curves were generated and statistical significance was assessed using the Gehan–Breslow–Wilcoxon test. For metabolic rate (CO₂ production) with more than two groups, data were first tested for normality. Datasets following a Gaussian distribution were analyzed using one-way ANOVA with parametric tests, whereas non-normally distributed datasets were analyzed using one-way ANOVA with non-parametric tests. For metabolic rate analysis between two groups, a two-tailed unpaired student t-test was used.

## Acknowledgements

We thank Véronique Tissot for providing patient imaging data and for her support of this study. We also thank the VDRC and Kyoto Stock Centers for providing *Drosophila* strains, as well as all members of the Schenck and Kleefstra labs for helpful discussions.

## Competing interests

The authors declare that they have no competing interests.

## Funding

This work was in part supported by a Vici grant from the Netherlands Organization for Health Research and Development (ZonMw, 09150181910022) to A.S, PSIDER grant from the Netherlands Organization for Health Research and Development (ZonMw, 10250022110003) to T.K. and H.B., ZonMw Open grant (09120012110034) to T.K. and J.D, and a Radboud Excellence Fellowship to S.G.J.

## Data and resource availability

All relevant data and resources can be found within the article and its supplementary information.

## Author contributions statement

Conceptualization: S.G.J., A.B., T.K., A.S.

Methodology: S.G.J., A.B., N.R., E.A.J.vG., E.H.J.G.A., J.D., J.G, H.B., S.M., D.G

Software: S.G.J., A.B., N.R., E.A.J.vG., J.D.

Formal analysis: S.G.J., A.B., N.R., I.MB., M.CT., E.A.J.vG., J.D.

Investigation: S.G.J., A.B., N.R., I.MB., F.K.

Data curation: S.G.J., A.B., N.R., E.A.J.vG., J.D.

Visualization: S.G.J., N.R., E.A.J.vG,

Supervision: S.G.J., M.CT., T.K., A.S.

Project administration: S.G.J., A.B., T.K., A.S.

Funding acquisition: S.G.J., H.B., J.D.,T.K., A.S.

Writing – original draft: S.G.J., A.B.

Writing – review & editing: All authors

## Notes

### Competing Interest Statement

The authors have declared no competing interest.

